# A multiscale approach for computing gated ligand binding from molecular dynamics and Brownian dynamics simulations

**DOI:** 10.1101/2021.06.22.449380

**Authors:** S. Kashif Sadiq, Abraham Muñiz Chicharro, Patrick Friedrich, Rebecca C. Wade

## Abstract

We develop an approach to characterise the effects of gating by a multi-conformation protein consisting of macrostate conformations that are either accessible or inaccessible to ligand binding. We first construct a Markov state model of the apo-protein from atomistic molecular dynamics simulations from which we identify macrostates and their conformations, compute their relative macrostate populations and interchange kinetics, and structurally characterise them in terms of ligand accessibility. We insert the calculated first-order rate constants for conformational transitions into a multi-state gating theory from which we derive a gating factor *γ* that quantifies the degree of conformational gating. Applied to HIV-1 protease, our approach yields a kinetic network of three accessible (semi-open, open and wide-open) and two inaccessible (closed and a newly identified, ‘parted’) macrostate conformations. The ‘parted’ conformation sterically partitions the active site, suggesting a possible role in product release. We find that the binding kinetics of drugs and drug-like inhibitors to HIV-1 protease falls in the slow gating regime. However, because *γ*=0.75, conformational gating only modestly slows ligand binding. Brownian dynamics simulations of the diffusional association of eight inhibitors to the protease - that have a wide range of experimental association constants (~10^4^ - 10^10^ M^−1^s^−1^) - yields gated rate constants in the range ~0.5-5.7 × 10^8^ M^−1^s^−1^. This indicates that, whereas the association rate of some inhibitors could be described by the model, for many inhibitors either subsequent conformational transitions or alternate binding mechanisms may be rate-limiting. For systems known to be modulated by conformational gating, the approach could be scaled computationally efficiently to screen association kinetics for a large number of ligands.

**Graphical TOC Entry:** 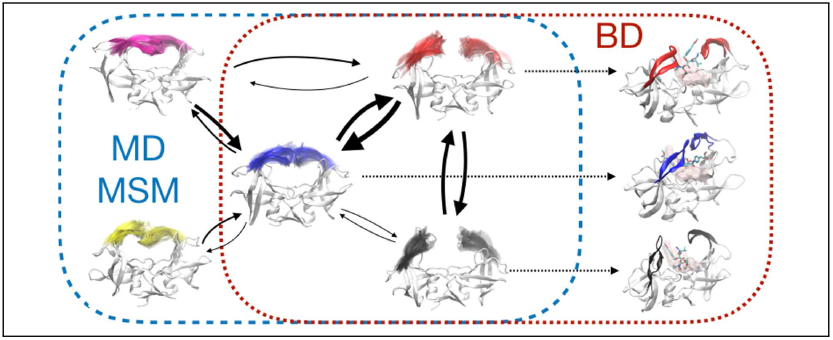

## 1 Introduction

Molecular association is central to virtually all biomolecular processes from regulation of metabolic networks and signal transduction to macromolecular assembly and drug efficacy in pharmacology. Extensive effort over several decades has led to the development of many computational methods to characterize molecular binding affinities.^1^ Increasingly, though, the role of binding kinetics, not just affinity, is becoming more widely appreciated in determining the behaviour of biomolecular interaction networks and especially *in vivo* drug efficacy.^2,3^

The binding of two molecules - such as a ligand to a protein - first requires them to encounter each other through a diffusional process, imposing an upper, diffusion-controlled limit - on the association rate.^4–6^ Factors such as steric specificity, hydrodynamic interactions and binding site accessibility can all cause reductions from this limit, whilst ligand-protein interaction energies can either enhance or further impede the association rate - for example, through electrostatic steering.^5,7–12^

Protein conformational changes can also modulate binding. A canonical view of these changes usually depicts two primary protein conformations in thermodynamic equilibrium, active and inactive, where the active conformation corresponds to the native ligand-bound state. ^13^ Both conformations are accessible to ligand binding. In a conformational selection (CS) mechanism, a ligand selects to bind only to the active conformation within the protein population ensemble. Induced fit (IF) is the opposite extreme, where the ligand first binds to the inactive conformation and then the protein is induced to transition into the active conformation by forming the required intermolecular contacts consistent with the native ligand-bound state. Both mechanisms have been shown to play a role in binding processes and their association kinetics.^14–16^ In general, however, both mechanisms can contribute to a given binding process. ^13,14^ Indeed, the relative contribution of IF versus CS increases with both ligand concentration^17,18^ and the rate of protein conformational transition.^18,19^ The existence of more molecular conformations can be described by a dock and coalesce (DC) mechanism^20,21^ in which simultaneous formation of all required native bound interactions upon or after initial protein-ligand binding is unlikely,^13^ thus precluding either purely CS or IF binding alone. This mechanism has been applied to the binding of highly disordered systems such as intrinsically disordered proteins (IDPs).^22–24^

Dynamic conformational changes in the protein can also alter binding site accessibility, introducing a gating step in the initial ligand binding process. ^25–27^ Gating theory for the interchange between two conformations, open and closed, ^27–31^ suggests the timescale of interchange (*τ_c_*) can play a significant role that depends on whether it is faster (*τ_c_* << *τ_d_*) or slower (*τ_c_* >> *τ_d_*) than the characteristic timescale of protein-ligand diffusional encounter (*τ_d_*), or in between. For fast gating, the ligand effectively always encounters the protein in an accessible (open) state - thus the gated (*k^g^*) and ungated (*k^ug^*) association rate constants are equal. For slow gating, *k^g^* = *γk^ug^*, where *γ*, the gating factor, represents the probability of finding the protein in a ligand-accessible state.

Brownian dynamics (BD) simulation methods offer a computationally inexpensive route for calculating ungated diffusional association kinetics, ^32^ in part due to frequent use of rigid bodies, simplified forcefields and implicit solvent models. They are well-established for protein-protein association rate constants^33,34^ and applied to protein-ligand systems.^35,36^ However, use of rigid structures poses challenges, especially becuase many biomolecular systems exhibit considerable flexibility and where CS, IF or DC mechanisms can be significant contributors to the binding kinetics. Rigid-body BD alone may therefore be applicable to mainly diffusional-limited binders. Multiscale methods that combine BD with molecular dynamics (MD) simulation methods may offer a possible solution to such limitations.^35^ A method, known as SEEKR, has been proposed where BD simulations are performed to reach the first milestone, followed by spawning of MD simulations at subsequent milestones. ^37–40^ Such methods have yet to be tested on protein-ligand systems with large conformational flexibility.

Furthermore, a plethora of methods have emerged that aim to compute ligand-receptor binding kinetics. ^41,42^ Approaches based on molecular dynamics (MD) simulations have become a focal point in this development process. ^43^ Unbiased high-throughput all-atom MD simulations coupled to Markov state models (MSMs) have proven a rigorous route to extract the kinetics and thermodynamics of biomolecular processes, as well as a detailed description of their respective pathways, including phenomena such as receptor-ligand, ^15,44–47^ and receptor-receptor binding,^48^ protein-folding,^49–52^ conformational changes^46,53–56^ and macromolecular assembly. ^57^

MSMs usually involve the projection of the high-dimensional conformational space of atomistic features into a lower-dimensional manifold of collective variables (CVs), followed by discretization of the resulting space into non-overlapping microstates. Assignment of the underlying MD simulation ensemble in terms of a sequence of these microstates allows interstate transition probabilities to be calculated at time intervals chosen to ensure a memoryless jump process, the condition of ‘Markovianity’.^58^ The resulting transition matrix, together with methods for further coarse-graining the MSM can then be used to probabilistically group together microstates into a smaller set of macrostates, that represent kinetically distinct interpretable conformations of the system as well as the corresponding coarse-grained equilibrium distribution and transition kinetics. ^59^

Advantageously, MSMs can account for detailed changes across complex degrees of freedom, such as ligand-receptor binding pathways and therefore can quantify the degree of CS, IF or DC involved. ^48^ Disadvantageously, they impose a huge computational demand in order to resolve such kinetics accurately. Estimation of association rate constants may require simulating up to ~10^3^-fold longer than the timescale of the binding process^3^ due to the dilute solute concentration in a typical MD simulation box. Furthermore, the choice of initial features, the process of dimensionality reduction to obtain CVs and clustering methods for discretisation can all affect the quality of the model. ^60^ Significant developments have been made in each of these aspects over the last decade, ^61^ including the derivation of a variational principle^62^ that enables selection of CVs to be systematized based on maximizing the decorrelation time of linear combinations of atomistic metrics, an approach known as time-independent component analysis, tICA.^63–65^ Furthermore, adaptive sampling techniques^66–68^ that explore the conformational space more efficiently and neural network approaches that learn optimal CVs^69^ can result in orders-of-magnitude reductions in required sampling.

Despite these major developments, application of MSMs remains prohibitive when scaled to multiple numbers of receptor-ligand systems, especially when typical experimentally measured associaton rate constants 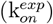 for many clinically relevant compounds are 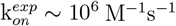 or slower.^3^ The high computational demand of unbiased MD has motivated a number of enhanced sampling methods that introduce biasing potentials and/or biased exploration, allowing faster exploration of rarely accessible regions of the conformational space. Recent developments in methods such as metadynamics, ^70,71^ bias-exchanged metadynamics, ^72^ Gaussian-accelerated molecular dynamics (GAMD),^73^ weighted ensemble sampling,^74^ scaled MD^75^ and adaptive multilevel splitting^76^ have been applied to rate constant calculations. Unfortunately, such methods either require predefined CVs - which are not always known - or the use of biasing potentials makes recovering the original kinetics challenging. ^77^

For the class of proteins that exhibit multiple conformational states that may gate ligand binding, we seek here to combine the advantages of MSMs for rigorously computing conformational kinetics with the advantages of BD approaches for rapidly computing ungated rate constants. HIV-1 protease is a suitably flexible, multi-conformation protein on which to develop such an approach and where the relative population, kinetic interchange and functional relevance of different conformations still remain incompletely understood as does the role of such conformations in gating ligand binding. The protease consists of a C_2_-symmetric homodimer with a pair of flexible *β*-hairpin structures, termed ‘flaps’ that gate access to an active site^78^ that structurally recognises^79,80^ and cleaves a number of sequence specific peptidic junctions in viral Gag and GagPol polyproteins. ^81–83^ Most crystal structures of HIV-1 protease are of ligand-bound complexes - such structures are invariably characterised by a ‘closed’ flap conformation^84,85^ where the flaps extend beyond but curl towards each other (Figure 1). A handful of apo-protein conformations exist. These include the well-characterised semi-open conformation^86^ differentiated by a reversal of flap handedness with respect to the symmetry axis of the dimer - with flaps extending beyond but curling away from each other (Figure 1). A spectrum of more open flap structures^87^ exist, marked by an absence of interflap interactions as well as the characteristic closed conformation. Structurally asymmetric conformations and flap dynamics have also been reported. ^88^

**Figure 1:**
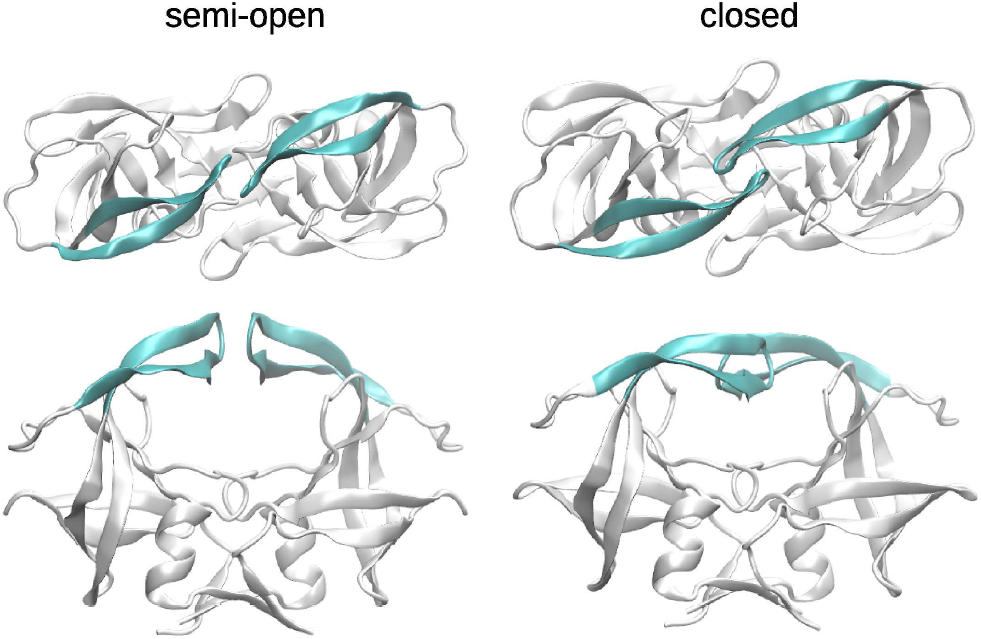
Characteristic conformations of the HIV-1 protease, illustrating a semi-open apostate (PDB:1HHP)^86^ and a closed state (PDB: 1KJ7),^80^ required for both substrate cleavage and observed for inhibitor binding. For clarity, the substrate is omitted from the closed state structure depicted here. The homodimeric protein is shown in ribbon representation with the gating flaps in cyan.

Natural substrate recognition, transition state stabilisation and peptide cleavage occur in the closed conformation. ^79^ Solution NMR experiments on the apo protein have reported both a fast (<10 ns) and slow (~100*μ*s) flap transition suggested to be flap tip curling and wide opening, respectively, as well as a dynamic equilibrium between three states of which the semi-open state is the dominant population.^89,90^ More recent NMR studies are inconsistent with this view, reporting a mixture of states where the closed conformation is favoured. ^91^ Although lateral binding or unbinding of inhibitors, small peptide substrates or auto-associating peptides to either the closed or semi-open conformations may be possible, ^53,72,92,93^ steric considerations require an open state to admit peptidic cleavage sites within Gag or GagPol polyproteins or natural subtrate analogs. MD simulations have also reversibly captured a mixture of closed, semi-open, open and wide-open states in the apo protein - open, characterised by the flaps curling back from semi-open to expose the active site and wide-open, by a significantly larger flap-tip separation^27,94–96^ - with a range of estimated timescales for flap opening (~10 ns), wide-opening (~500 ns) and closing (~50 ns) from a semi-open state.

BD simulations in a coarse-grained model of a two state open-closed system suggested a slow gating regime and showed good agreement with experimental rate constants for a number of inhibitors in the 10^5^-10^6^ range.^27^ However, surface plasmon resonance (SPR) measurements association rate constants for a large array of inhibitors of HIV-1 protease, including the 9 FDA-approved drugs in clinical use,^97^ showed values ranging widely from 10^2^-10^10^ M^−1^s^−1^. This suggests that a significant degree of CS, IF or mixed DC meachnism may be at work during ligand binding and possibly in a ligand-dependent way. Furthermore, as more than two states exist and the open state is an intermediate state, then gated binding to an open state would still only be the first step of the process. A second step, requiring some degree of structural rearrangement is subsequently necessary to arrive at the final bound closed state. Simulations have suggested this second step may be rate limiting^98^ and support the idea that different ligands may vary in their preferred binding mechanism. ^99,100^

Here, using HIV-1 protease as a model system, we report a novel method that combines BD and MSMs built from all-atom MD simulations to calculate the extent of conformation gated ligand binding. This is done by combining the two approaches within a theoretical multi-state gating framework. We build on previous gating theory^25–27^ to determine a gating factor in terms of the accessible and inaccessible ensemble of conformational macrostates of a MSM. Next we reconstruct the full apo-protein conformational kinetics of HIV-1 PR from a MSM based on all-atom MD simulations, identify accessible and inaccessible states - including a previously unidentified state - from which we obtain the resulting gating factor. Finally, we compute the ungated association kinetics for several ligands binding to each of the set of determined accessible states using BD simulations, applying the theoretical framework to arrive at final gated rate constants. This results in fast and scalable calculations of absolute values of protein-ligand biomolecular association rate constants that approach experimental values, where gating is the dominant factor in limiting binding and allows us, in some cases, to dissect various binding mechanisms.

## Theory

We build on the theoretical formalism of MSMs^101^ and gating theory,^25–27^ in order to derive a term for the required gating factor to modulate ligand binding association kinetics in terms of the transition kinetics between conformational macrostates exhibited by the unbound target receptor.

Let us consider a Markov state model (MSM) of a stochastic process {*X_n_*} which is sufficiently described by a transition matrix **T**. The elements of the matrix are the conditional probabilities

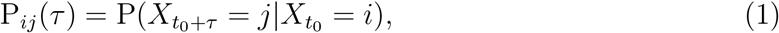

describing the probability of a transition from a current state *i* to the next state *j* within the lag time *τ*. A MSM fulfilling the Chapman-Kolomogrov equation **T**(*n* · *τ*) = **T**(*τ*)^*n*^, in other words a vector holding the description of the state distribution at any time *t* = *t*_0_ + *nτ*, can be calculated from an initial distribution **p**(*t*_0_) by

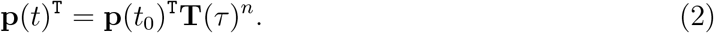

After sufficient time the probability distribution approaches the stationary distribution *π* = **p**(*t* → ∞), which contains the probabilities of the single states in equilibrium. Further application of the transition matrix on *π* has no effect

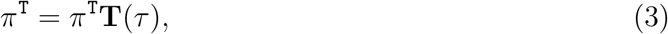

which forms an eigenvector equation. When such a stationary distribution exists a property, the detailed balance, can be derived from Eq.3 and 1.

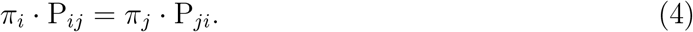

This property is physically important and describes the reversibility of the MSM.

The MSM transition matrix is estimated from the observed/counted transitions between the states in the single MD simulations. An important parameter for the accuracy of the estimation is the lag time *τ*. Ideally, the eigenvalues λ_*i*_ of the MSM should be independent of *τ*. Therefore *τ* should be chosen such that the implied relaxation time scales are approximately constant

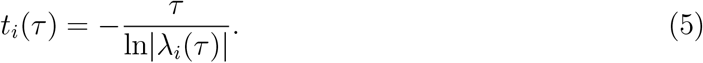

To obtain those *t_i_* for a MSM, the lag time might easily exceed the given time scales and therefore our interest lies only in the largest *u* eigenvalues λ_*i*_(*τ*), which represent the slowest state transitions. For projected MSMs built on the reduced set of slow-transition eigenvalues, Eq.2 does not in general hold anymore. Rather, in order to coarse-grain an MSM into a meaningful and smaller set of macrostates, a hidden markov model (HMM) can be built to describe u such states at any time step *kτ*.

The HMM can then be expressed as:

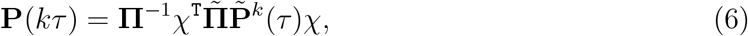

where 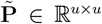 is a transition matrix of *u* kinetic important coarse-grained states and 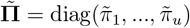 contains the corresponding stationary distribution of the coarse-grained u-state HMM. *χ* is a matrix, whose elements *χ_Ij_* ∈ ℝ^*u*×*v*^ hold the probability to observe a macrostate *I*, while the system is in microstate *j*. **Π** = diag(*π*_1_,...,*π_v_*) is the stationary distribution of the *v*-state MSM.

To calculate a coarse-grained matrix, the microstates are grouped into a membership matrix **B** ∈ ℝ^*v*×*u*^, the elements of which contain the probability that a microstate *i* participates in a macrostate *J* and are computed by linear transformation of the first *u* eigenvectors λ_*i*_ (PCCA+ and PCCA++ methods). χ is then calculated as

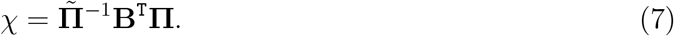

Also the *u* × *u* transition matrix of Eq.6 can be calculated from the membership matrix **B** as

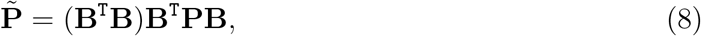

where **P** is the *v* × *v* MSM transition matrix. Together with Eq.7, Eq.8 is used to estimate the HMM.

A transition rate matrix **K** can then be derived, which describes the state transitions in terms of the actual kinetics. The relation between the rate and probability matrix is given by the equation

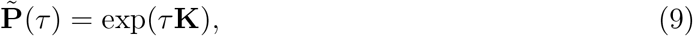

where *τ* is here again the lag time at which the given HMM is constructed.

Let us now consider a multistate receptor that permits significant ligand access to only a single accessible state *r*, as described previously.^27^ In a regime where ligand binding is limited by conformational gating to *r* and not by a subsequent induced fit association rate constant (*k^if^*), the overall bimolecular association rate constant (*k*_on_) is dominated by the gated association rate constant (*k*^g^) and can be expressed as:

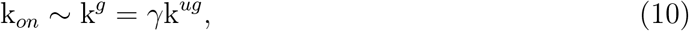

where *γ*=k_*acc*_/(k_*acc*_+k_*nacc*_), is the conformational gating factor and represents the probability of the ligand finding the apo-receptor in state *r* (Figure 2). The first-order rate constants k_*acc*_=Σ_*i*≠*r*_ k_*ir*_ and k_*nacc*_=∑_*j*=*r*_ k_*rj*_ describe the overall transition kinetics between accessible state *r* and all other non-accessible states in the apo-receptor. Here, *k_ij_* represent the (off-diagonal) elements of the transition rate matrix **K**, representing the net flux from a single metastable state *i* to another metastable state *j*, and which can be computed directly from the HMM of the apo-receptor, as described above.

**Figure 2:**
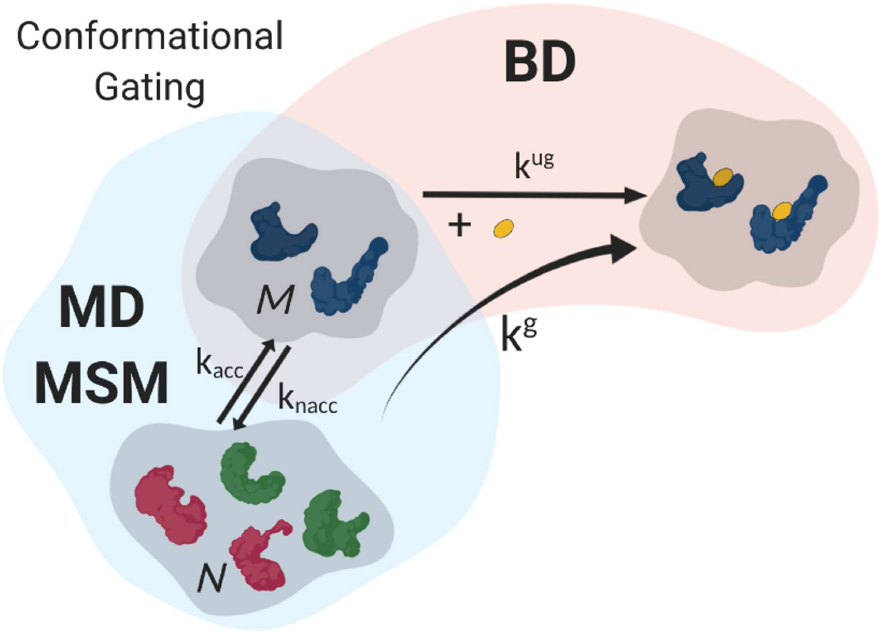
Schematic of coupling approach between BD and MD/MSM to compute gated ligand binding to accessible states of a multiconformation receptor. A conformational gating factor (*γ*), representing the probability of accessing the accessible states is derived from the previously described hidden Markov model (HMM) transition matrix of the apo-state conformational kinetics (see Theory), where *γ*=k_*acc*_/(k_*acc*_+k_*nacc*_) and where k_*acc*_ and k_*nacc*_ are the first-order rate constants for transitioning into the accessible and non-accessible states, respectively. BD simulations are then performed for binding to the accessible states yielding a second-order ungated association rate constant (k^*ug*^). For ligand-receptor complexes in which other binding mechanisms are not rate-limiting, the overall association kinetics can be approximated by a gated rate constant, k^*g*^=*γ*k^*ug*^.

We now expand this model more generally to the case of multiple accessible states. The apo-state space is divided into two disjoint subsets, containing all non-accessible *N* and all accessible *M* states. The first-order conformational rate constants between the two sets of states can then be expressed as the sum of transition rates from non-accessible *n* ∈ *N* to accessible *m* ∈ *M* states such that k_*acc*_ Σ k_*nm*_ and vice versa k_*nacc*_ Σ k_*mn*_. A gating factor (*γ*) can then be defined for the overall transition of the accessible states:

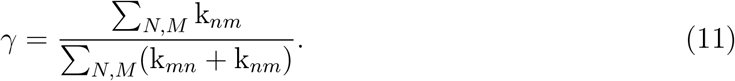

However, *γ* alone does not contain the information about the respective contributions of the accessible states. To account for these we define the state specific gating factor Γ_*m*_ = *γϵ_m_*, where *ϵ_m_* represents the relative population of state *m* ∈ *M* and which can also be computed from the HMM. By combining with ungated rate constants 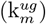 computed from BD simulations for each state *m*, the overall gated association rate constant (k^*g*^) for the set of states, *M*, is then given by:

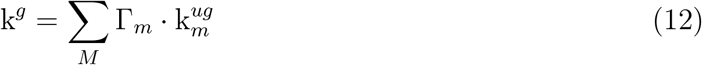

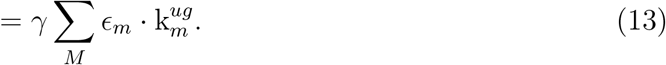

With this formalism, the combined ungated rate constant (k^*ug*^) for the set of states, *M*, is then given by:

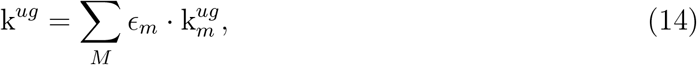

and in the limiting case where there is only one accessible metastable state, *ϵ_m_* = 1, we recover the single-state gating factor. Extending the previous formalism^27^ to multiple accessible states, ‘slow’ and ‘fast’ gating regimes can be differentiated in relation to the characteristic diffusion time 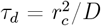, where *r_c_* is the collision distance between the molecules and *D* is the relative translational diffusion constant by:

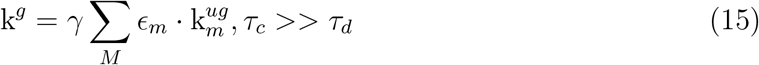

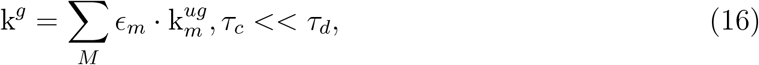

where *τ_c_* is the characteristic timescale of conformational gating given by:

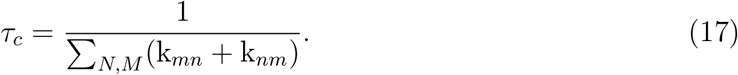

## Materials and Methods

### Initial preparation

Atomic coordinates for wildtype dimeric HIV-1 protease in the semi-open conformation were extracted from the 1HHP crystal structure^86^ in the Protein Data Bank.^102^ Those for a wide-open conformation were taken from the reported structure in our previous study. ^95^ The standard AMBER forcefield (ff03)^103^ was used to describe all protease parameters. A dianionic state was assigned to the catalytic dyad as, at physiological pH, the unbound protease is unprotonated, whilst mono-protonation is thought to occur upon or after ligand binding.^104–107^ The VMD software package^108^ and the LEAP module of the AMBER 16 software package^109^ were then used to build the initial systems. Each system was solvated using atomistic TIP3P water^110^ in a cubic box extending at least 7 Å around the complex and then electrically neutralised with an ionic concentration of 0.15 M NaCl, resulting in fully atomistic systems for both semi-open and wide-open conformations of ~40,000 atoms.

### Equilibration and production MD simulations

All equilibration and production simulations were performed using ACEMD. ^111^ During equilibration the positions of the heavy protein atoms were restrained by a 10 kcal/mol/Å^2^ spring constant and conjugate-gradient minimisation was performed for 2000 steps. The hydrogen atoms and water molecules were then unrestrained and allowed to evolve for a total of 500 ps at 300 K in the NVT ensemble to ensure thorough solvation of the complex and to prevent premature flap collapse. ^112^ The magnitude of the restraining spring constant was set to 1 kcal/mol/Å^2^ for the following 500 ps, then to 0.05 kcal/mol/Å^2^ for 500 ps, then finally to zero for 500 ps. The temperature was maintained at 300 K using a Langevin thermostat with a low damping constant of 0.1/ps and the pressure was maintained at 1 atm using a Nose-Hoover barostat. The sytem was finally equilibrated for 6 ns of unrestrained simulation in the isothermal-isobaric ensemble (NPT). The long range Coulomb interaction was handled using a GPU implemention^113^ of the particle mesh Ewald summation method (PME)^114^ with grid size 84×84×84. A non-bonded cut-off distance of 9 Å was used with a switching distance of 7.5 Å. For the equilibration runs, the SHAKE algorithm^115^ was employed on all atoms covalently bonded to a hydrogen atom with an integration timestep of 2 fs. Production simulations took place in the NVT ensemble. The hydrogen mass repartitioning scheme^116^ was used, allowing a timestep of 4 fs. All other parameters were kept the same as in the equilibration phase. The production data set consisted of 465 and 499 simulations started from semi-open and wide-open conformations respectively, each for 130 ns with coordinate generation every 100 ps.

Our molecular simulation protocol for the HIV-1 protease has been previously validated using NMR S^2^ order parameters. ^95^

### Determination of collective variable space

Before MSMs can be constructed from MD simulation data, the high-dimensional configuration space of atomistic coordinates needs to be projected onto a low-dimensional, but meaningful, collective-variable (CV) space. Finding the appropriate CV space is still a major challenge, despite considerable efforts at systemisation. ^63–65^ A minimum fluctuation alignment (MFA) method, introduced previously^53^ and developed further here, was used to determine a set of information-rich collective variables that describe flap conformations in HIV-1 PR.

Minimum fluctuating (MF) residues for the protease were identified as residues 23, 24, 85 and 89-92 of each monomer. The homodimeric C_2_ symmetry of the protease makes identical MF residue subsets on both monomers a sensible choice of vector termini. The x-axis was thus chosen to be the vector between the center of mass (COM) of the backbone atoms of residues 89-92 of the second and first monomers respectively. A second vector was selected (between the COM of residues 23,24 and 85 of each monomer respectively). The resultant z-axis therefore approximated the C_2_ symmetry axis. The selected residues are either within the *α*-helix of the protease or within the *β*-sheet region that supports each side of the activesite and thus exhibit minimal RMSFs compared to other residues.^95^ Due to the choice of the x-axis, the resultant y-axis was similar to the vector describing the binding conduit of bound peptides in the active site. ^79,80^

A 3D metric based on the re-aligned cylindrical polar components of the I50-I50′ C_*α*_-atom flap tip separation vector, λ(λ_*r*_, λ_⊖_, *λ_z_*), was then devised. A wrapping angle of 50° was added to all values of λ_⊖_ with a wrapping modulus of 360°. This extends the original 1D Cartesian λ_*x*_ metric used earlier^53^ and was employed as a set of collective variables to describe the conformational transitions of the system.

### Markov state model

The software package pyEMMA2 ^101^ was used for Markov state model construction and analysis. The 3D λ-space was used as the set of collective variables in which to construct the models. The data were subsequently clustered into 100 microstates using k-means clustering in this 3D CV-space. Discretized state labels were assigned and Markov state models (MSMs) with Bayesian error estimation were built at various lag-times (*τ*) and the computed relaxation timescales analyzed as a function of *τ*. Constant relaxation timescales for the slowest transitions were exhibited at *τ*= 30 ns, significantly shorter than the length of all trajectories used, with time scale separation suggesting a set of five kinetically distinct macrostates (C1-C5). A subsequent MSM was then estimated using a lag time of 30 ns. The 100-state MSM at *τ*= 30 ns was coarse-grained into a five-state hidden Markov model (HMM) and validated using a generalized Chapman-Kolmogorov test. Fuzzy clustering of microstates into macrostates was performed and the resulting membership (**B**) and observation (χ) distribution matrices computed. The transition matrix 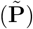 between the smaller set of five meaningful macrostates (C1-C5) was then computed using Equations 7 and 8. The stationary distribution of the five states and the transition kinetics between these states was then extracted from the HMM.

### Structural characterization

Representative snapshots of the kinetically distinct macrostates (C1-C5) were extracted for structural characterization. Macrostate C1 was noted to be an antisymmetric doublet-state (see Results), thus representative structures were extracted separately for each of the corresponding substates C1a and C1b. For each macrostate, the microstate with the largest observation probability was selected. Within each corresponding microstate, conformers were ranked in order of proximity in the λ-space to the centroid and the closest 1000 conformers extracted for analysis. Microstates for C1a and C1b corresponded to those with the first and second highest observation probability, respectively, for macrostate C1. In addition, various structural features from the extracted ensembles were compared with respect to the first replica of a previous ensemble MD simulation of the HIV-1 protease in a closed conformation and bound to the SP1-NC natural octapeptide.^117,118^ Analysis was implemented using Python scripts based on the MDAnalysis toolkit. ^119^ Distributions of the root-mean-squared-deviations (RMSDs) of the C_*α*_ backbone with respect to semi-open (1HHP) and closed (1KJ7) reference crystal structures were computed as were the distributions of the absolute flap-tip separation (λ). Instantaneous hydrogen bonds were computed based on a donor-acceptor distance threshold of ≤3.5 Å and a donor-hydrogen-acceptor angle of ≥150°. Only those hydrogen bonds with an exhibition frequency *f_hb_* ≥ 0.5 were considered stable - the corresponding mean distances and angles were computed from the exhibited snapshots. Hydrophobic contacts were determined from sidechain-sidechain distance analysis; the minimum non-hydrogen atom distance between pairs of sidechains was computed for each conformer. A hydrophobic contact was assigned to hydrophobic residue pairs with a mean minimum distance ≤ 5 Å across the corresponding ensemble/trajectory. Volumetric maps of the averaged weighted atomic densities for each apo-conformer ensemble and the natural octapeptide from a former study^117^ were computed with the Volmap tool in VMD^108^ using a grid spacing of 0.5 Å.

### Brownian dynamics simulations

Atomistic coordinates for the 100 conformers most proximal to the microstate centroid in each structural ensemble were extracted to use in Brownian dynamics (BD) simulations to compute diffusional association rate constants for a set of inhibitors with experimentally determined rate constants.^97^ Protein partial atomic charges were assigned using the ff14SB forcefield. ^120^ Atomistic coordinates and partial charges for the inhibitors were extracted from a previous study.^121^ PQR files were generated using the PDB2PQR module in AMBER18.^122^ Electrostatic potential grids for the protein structures and for the inhibitors were computed using APBS^123^ with a grid spacing of 1 Å, an ionic concentration of 150 mM and a temperature of 298K, corresponding to experimental conditions, as well as a solvent dielectric constant of 78, a solute dielectric constant of 2 and an ionic radius of 1.5 Å. For the calculation of the electrostatic potential, the linearized Poisson-Boltzmann equation was solved under the single Debye-Hückel dielectric boundary condition. All BD simulations were carried out using SDA7.2.3,^124^ and association rates constants were computed using the Northrup-Allison-McCammon algorithm. ^32^ SDA employs an effective charge model (ECM), ^125^ where the charge on a reduced number of centers is fitted to reproduce, in a uniform dielectric medium, the potential computed with APBS for a heterogeneous dielectric medium in the region around the molecule. ECM charges for the protein were determined using the standard protocol in SDA, those for the ligands were determined from a newly developed protocol. ^126^ Electrostatic and hydrophobic desolvation grids were generated with a grid spacing of 1 Å. The ionic strength was set to zero for computing the electrostatic grid, with electrostatic and hydrophobic desolvation factors of 0.36 and −0.013 assigned, respectively. The solute and probe radii were both set to 1.4 Å.

Translational and rotational diffusion coefficients were calculated for all solutes using Hydropro^127^ with an atomic element radius (AER) of 2.9 Å and 1.2 Å for protein and ligands respectively, with a *σ_min_* of 1.0 Å and a *σ_max_* of 2.0 Å. For each macrostate-ligand combination, a diffusional association rate constant was computed from a set of 100000 trajectories (100 protein conformers × 1000 replicas). Each trajectory was started with an initial center-to-center distance of 150 Å between the solutes and stopped when the solutes reached a seperation of 250 Å. A variable time step was used, linearly ranging from 1 ps at a solute-solute separation of 50 Å to 20 ps at a separation of 90 Å. A distance threshold of 3 Å was set for pairs of contacts to be assigned as independent reaction criteria that described formation of a receptor-ligand encounter complex. Reaction criteria included polar interactions at 3.5 Å defined in terms of acceptor-donor pairs, as well as methyl, halogen and aromatic interactions at 4.5 Å, defined in terms of *π-π* pairs from the above-mentioned crystal structures of the corresponding complexes, filtering out any residues that did not match the sequence of the apo-protein (PDB: 1HHP) and including symmetrically equivalent contacts due to the C_2_ symmetry of the protease dimer. Reaction criteria coordinates were taken for each BD simulation after superimposition of the C_*α*_ backbone of residues 1-43 and 58-99 in each monomer of a given apo-conformer with the corresponding original inhibitor-bound complex. An encounter complex was counted whenever a sufficient number of contacts from the subset of pre-defined reaction criteria was observed within a specified distance. The association rate constant was initially calculated using a range of definitions of encounter complex in terms of number of contacts, {1,...,5} and their corresponding distances from a minimum of 3 Å and with increasing window intervals of 0.5 Å up to 20 Å. Ungated association rate constants were subsequently computed by requiring the formation of 3 independent contacts shorter than 4 Å.

BD docking simulations were performed for each of the ligands diffusing to each of the protein macrostates under the same reaction criteria as for the association rate calculations to check if the reaction criteria corresponded to ligands binding within the active site of the protease. 10000 trajectories were run for protein conformers with the highest number of encounters obtained in the association rate constant calculation simulations, and complexes were recorded at a 4 Å reaction criterion window distance.

## Results

Ensemble molecular dynamics simulations resulted in ~1.25 ×10^6^ molecular conformers for subsequent analysis. In order to reveal and characterize meaningful conformational states from this ensemble, we constructed a converged Markov state model (MSM) based on a suitable projection of the atomistic coordinates into a three-dimensional collective variable space. This enabled us to identify a small set of discretised macrostates of the system, compute their equilibrium distribution and transition kinetics, and then analyse representative constituent conformers belonging to each macrostate to reveal distinct structural features. This in turn allowed the differentiation of the macrostates that were accessible and non-accessible to ligand binding and the subsequent computation of a gating factor. Brownian dynamics simulations enabled computation of the ungated association rate constant to accessible states from which the overall gated rate constant could be computed for a set of ligands.

### Kinetic network of HIV-1 protease conformations

The 3D λ-state-space in cylindrical polar coordinates (*λ_r_*, λ_⊖_, λ_*z*_) was chosen as a collective variable space in which to construct Markov state models (MSMs). The λ-state-space was discretised into 100 microstates. MSMs constructed using Bayesian error estimation across a range of lag-times (*τ*) show independence of the relaxation timescale (*t_n_*) for the slowest-order transitions at *τ* = 30 ns (Figure 3(a)) and a significant timescale separation between the fourth and fifth non-stationary eigenvectors with t_4_/t_5_ > 2 (Figure 3(b)).

**Figure 3:**
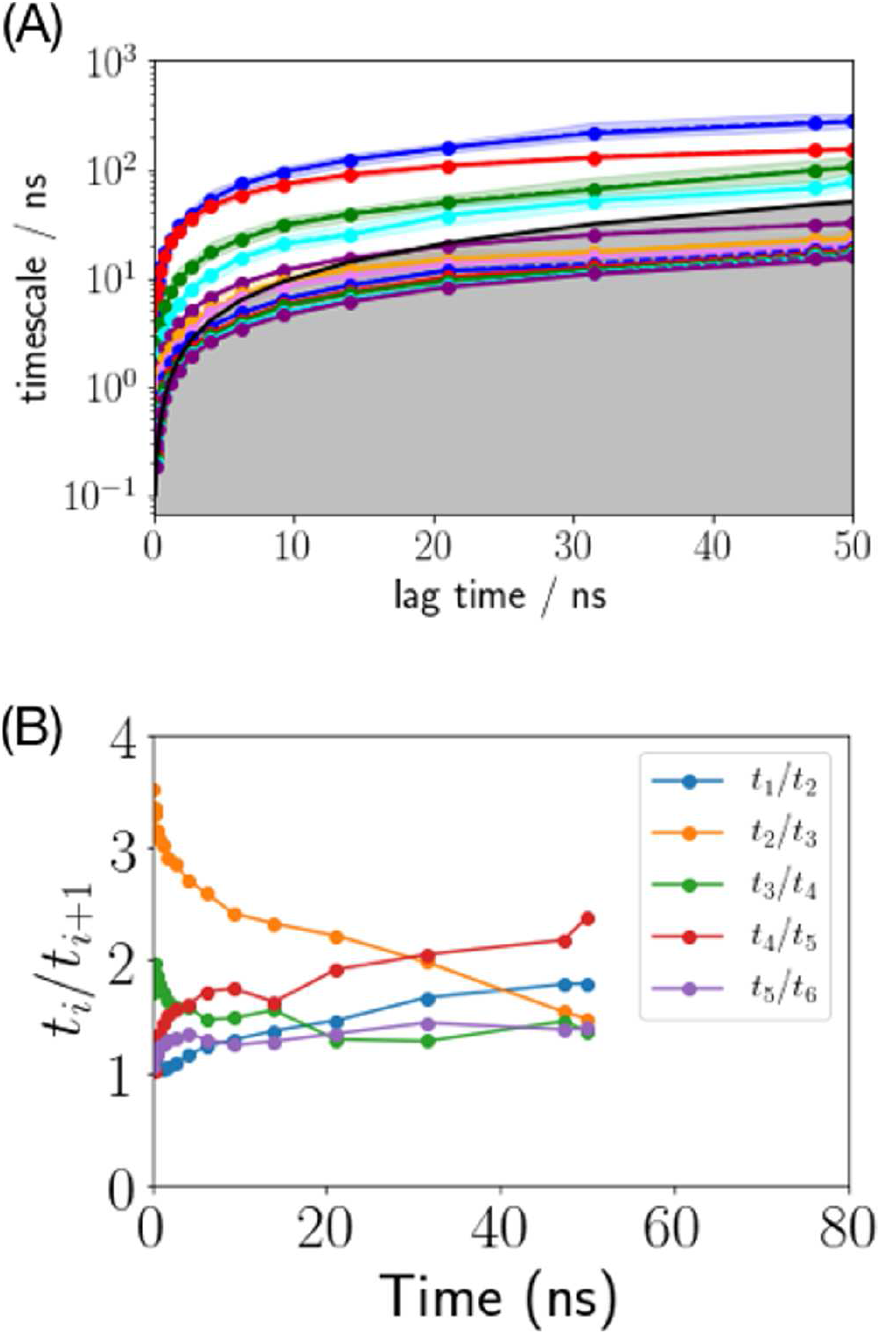
(a) Relaxation timescales (t_*n*_) for the dominant non-stationary eignvectors of the MSMs constructed in the λ-space at increasing lag times (*τ*) with Bayesian error estimation. Independence of the relaxation timescale with lag time is observed at *τ* = 30 ns. (b) Plot of the ratio of proximal relaxation timescales (t_*i*_/t_*i*+1_) shows a signifcant timescale separation of t_4_/t_5_ > 2 at *τ* = 30 ns suggesting the existence of five kinetically meaningful metastable states.

Consequently a MSM was constructed with Bayesian error estimation using 100 microstates and τ = 30 ns and the microstate equilibrium distribution computed. A potential of mean force (PMF) landscape, computed for 2D projections of the λ-state-space and weighted by this equilibrium distribution (Figure 4(A)), samples a large region, ranging from 0 < λ_*r*_ < 40 Å, 0 < λ_⊖_ <360° and −15 < λ_*z*_ < 15 Å. This PMF differentiates several significant energetic minima with λ(λ_*r*_, λ_⊖_, λ_*z*_) centers at λ ~(8 Å,40°,±5 Å), λ ~(5 Å,100°,0 Å), λ ~(20 Å,150°,0 Å) and λ ~(8 Å,300°,0 Å), with minima values of −0.5, −4.5, −3.5 and −2.5 kcal/mol respectively, as well as one saddle point with a center at λ ~(10 Å,120°, 0 Å) and an energy of −2.0 kcal/mol.

**Figure 4:**
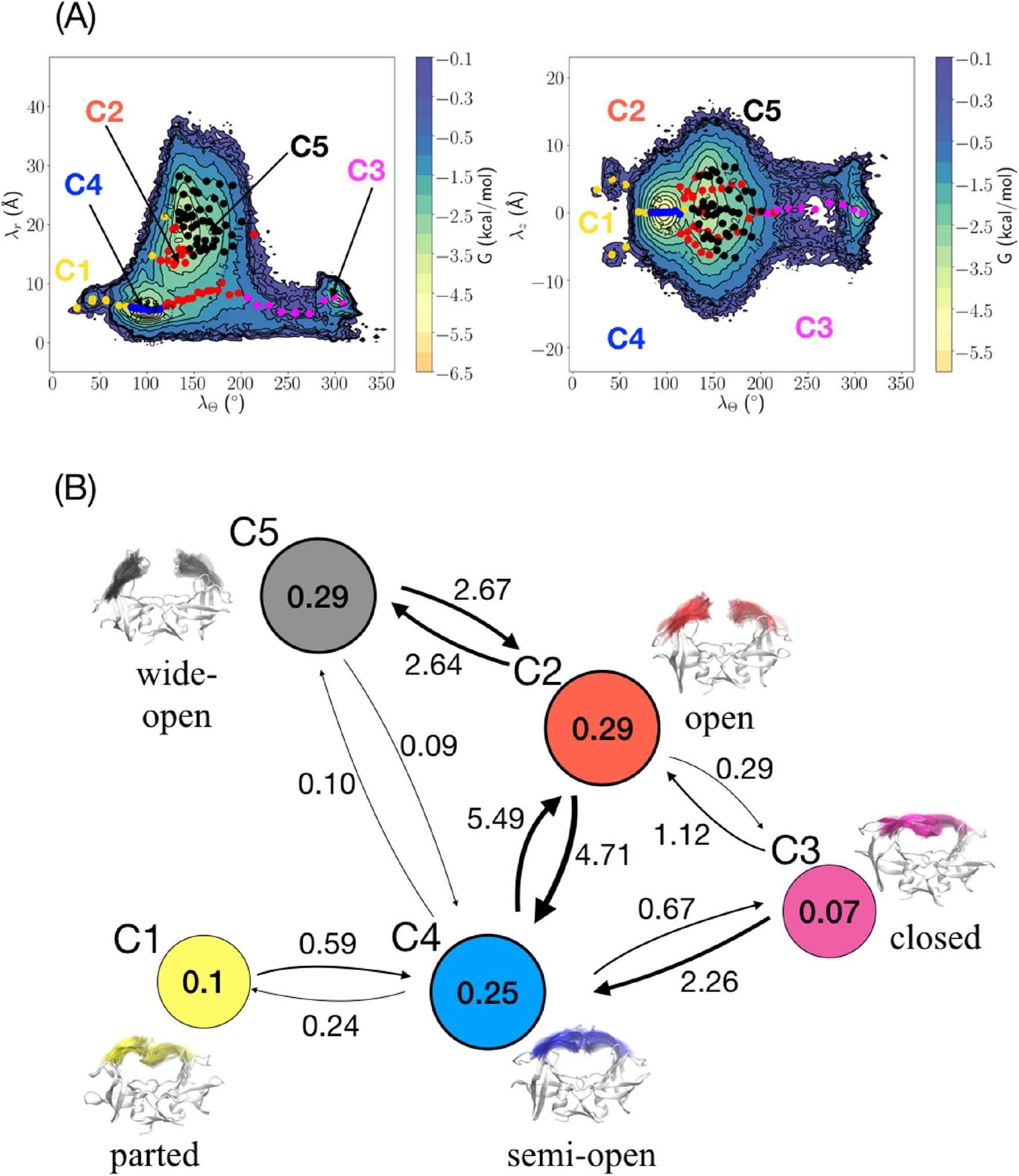
(A) Potential of mean force (PMF) of HIV-1 PR in the apo-state in the λ(λ_r_, λ_Θ_,λ_*z*_)- collective variable space, projected onto λ_*r*_-λ_Θ_ and λ_*z*_-λ_*z*_. Values are in kcal/mol. Energetic minima correspond to frequently sampled regions of the λ-space weighted by the equilibrium distribution of discretised microstates of the MSM. A HMM based on five macrostates (C1-C5) has clusters of corresponding microstates whose λ-centroids are overlayed on the PMF. (B) Kinetic network between the five coarse-grained macrostates, corresponding to conformations C1-C5, determined by the hidden Markov model (HMM). Stationary probabilities are shown in the circles, forward and reverse first order transition kinetics are shown by arrows with values in *μ*s^−1^. Displayed structures for each macrostate are superpositions of conformers belonging to the most probable microstate given the corresponding macrostate.

Timescale separation analysis implies that five kinetically meaningful macrostates exist, with the two slowest transitions in the system being around or larger than the 100 ns-timescale. Based on this a corresponding five-state coarse-grained hidden Markov model (HMM) was computed. This further clustered the microstates into kinetically separated macrostates, C1-C5 (Figure 4(A)). This model was validated by performing a generalized Chapman-Kolmogorov (C-K) test (Supporting Figure S1(A)). The C-K test showed good agreement with the predicted probability decay from the model at *τ* = 30 ns 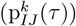 and the expected decay from models constructed at multiple (*k*) lag times (p_*Ij*_(*kτ*)). Moreover, the probabilities, χ, of observing each of the 100 microstates across the five macrostates (see Theory) were computed and projected onto the λ-space (Supporting Figure S1(B)). This analysis confirmed that the five macrostates also exhibited mutual exclusivity of microstate probabilities and therefore could be interpreted as distinct states. The microstates that constitute most macrostates correspond well with the energetic minima in the PMF. The only exception is C2, whose microstates seem to be primarily distributed in a transitional region between C4 and C5.

The HMM yielded a kinetic network of the stationary distributions and their respective first-order transition kinetics (Figure 4(B)). The five states, C1-C5, exhibit different stationary probabilities (*ρ*) with *ρ*_1_ = 0.1, *ρ*_2_ = 0.29, *ρ*_3_ = 0.07, *ρ*_4_ = 0.25 and *ρ_5_* = 0.29 corresponding to zero-centered Boltzmann-weighted free energies of ~-0.2, −0.8, 0, −0.7 and −0.8 kcal/mol, respectively. Thus the free energy differences betwen all five states are thermodynamically barely indistinguishable from thermal noise (~0.6 kcal/mol).

Nonetheless, there are significant differences in the transition kinetics between the various states. Only direct fluxes faster than 100 *μ*s are shown (≥ 3×10^−4^/*τ* = 0.01 *μ*s^−1^). Forward and reverse transitions between C2 and C5, as well as between C2 and C4, are well within the sub-*μ*s timescale (> 1 *μ*s^−1^), although the rate for direct transition between C4 and C5 is ~0.1 *μ*s^−1^. The C1 conformation only directly accesses the C4 conformation, and does so with kinetics slower than the *μ*s-timescale, whilst the C3 conformation accesses both C2 and C4 conformations with forward and reverse kinetics faster and slower than 1 *μ*s^−1^, respectively. Only the C4 conformation directly accesses all other conformations on a timescale faster than 100 *μ*s.

### Structural characterization of principal conformations

The conformational features of representative structures corresponding to each of the macrostates were analyzed. Probabilistic clustering of microstates into macrostates via the observation probability matrix (*χ_I,j_*) enables selection of the most likely microstates that represent each macrostate (Supporting Table S1). For each macrostate, the most probable microstate largely coincides with the corresponding minimum or minima in the λ-space (Supporting Figure S2). Macrostate C1 is characterized by two distinct minima in the λ-space equidistantly separated from the origin along λ-z. This implies a symmetry-related doublet state. Moreover the two most probable microstates for C1 correspond to each of these minima - thus C1 was split into two sub-states, C1a and C1b, in subsequent analyses corresponding to the +λ_*z*_ and -λ_*z*_ regions respectively. A set of 1000 closest conformers was thus extracted from the most probable microstate of each macrostate based on the ranked proximity of each snapshot to the corresponding microstate centroid in the 3D cylindrical polar λ-space.

Visual inspection of each set of conformers reveals that several macrostates correspond to known conformations of HIV-1 protease (Figure 4(B)), as well as a previously unidentified conformation. Hydrogen bonds and sidechain-sidechain contacts between each flap and the opposite (*trans*) monomer were analyzed (Supporting Tables S2 and S3). C2 and C5 correspond to open and wide-open conformations, respectively, where the flaps are pulled back enabling sterically unhindered access to the active site (Figure 4(B)). These two conformations exhibit no trans flap-monomer interactions. Macrostates C3 and C4 correspond to the characteristic closed and semi-open conformations, respectively, where the flaps extend over the active site but with opposite handedness (Figure 5 and Figure 6(A)-(B)). In C3, the I50 sidechain interacts with the characteristic array of hydrophobic residues (V32′, I47′, I54′, I80′ and I84′) on the opposite monomer and symmetrically vice versa. In C4, these contacts are missing, and are instead replaced predominantly a smaller set of hydrophobic contacts (I50-I50′,I50-F53′ and F53-I50′) and a symmetric pair of inter-glycine amide hydrogen bonds (G51–N-H–O-G49′ and G5F–N-H–O-G49). C1a and C1b are identified as anti-symmetric sub-states of a novel conformation, where one flap is extended as in the semi-open conformation whilst the other orients under it and traverses across the active site towards the other monomer (Figure 5). They share a subset of hydrophobic contacts with both C3 (C1a: I50-V32′ and I50-I47′, C1b: V32-I50′ and I47-I50′) and C4 (C1a: F53-I50′, C1b: I50-F53′) but also exhibit sidechain contacts (C1a: I50-D30′ and M46-I50′, C1b: D30-I50′ and I50-M46′) and a backbone hydrogen bond (C1a: G48′-N-H–O-I50, C1b: G48-N-H–O-I50′) that are unique to them and which serve to stabilise these conformations. In particular, the I50-D30′/D30-I50′ contacts bring the corresponding flap tip diagonally across to the opposite side of the active site.

**Figure 5:**
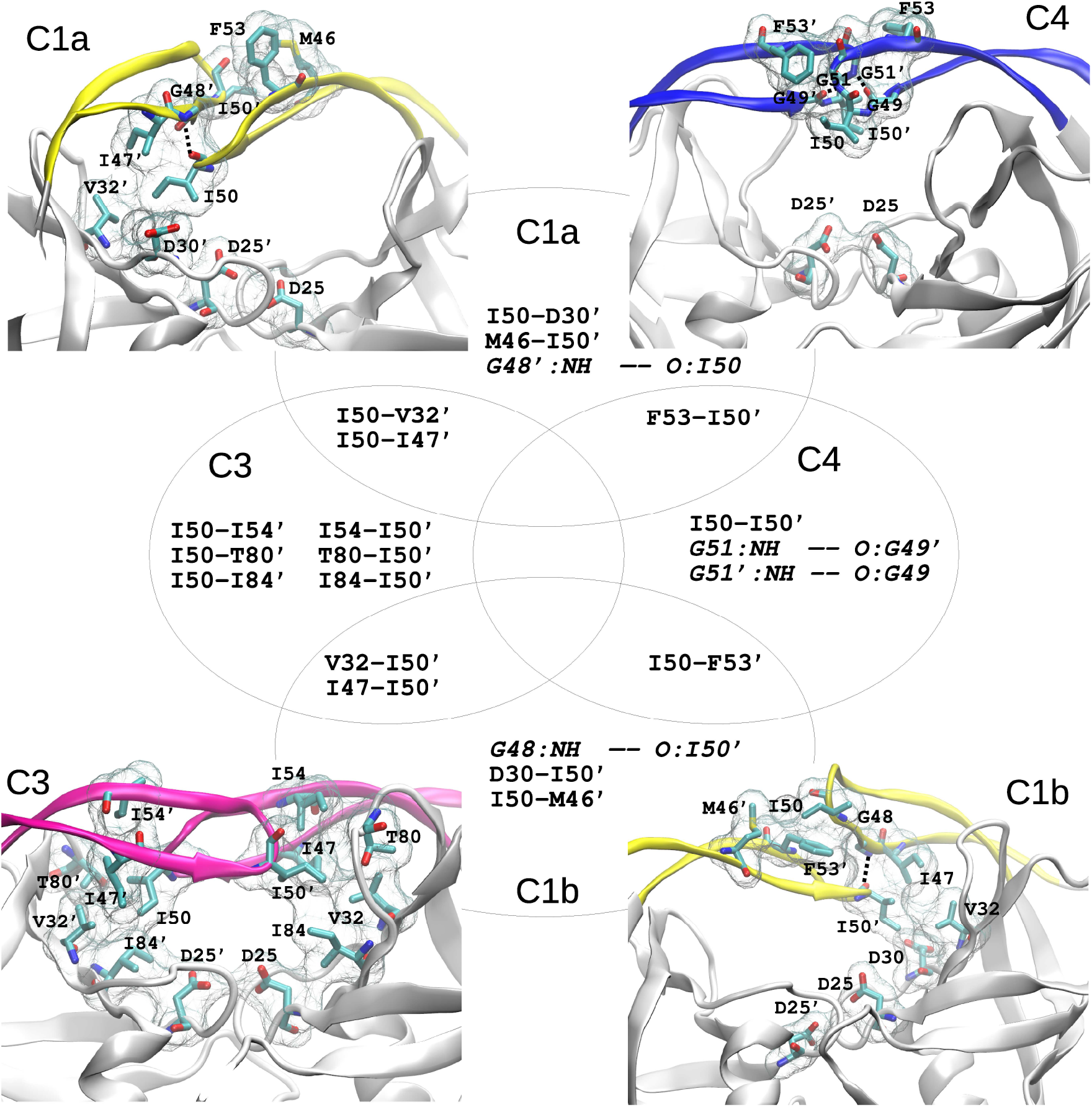
Mututally exclusive and shared sidechain-sidechain and hydrogen bond interactions between each flap region and the alternate monomer across different macrostates. C2 and C5 do not form any such interactions and are not displayed. Hydrogen bonds are marked with dashed black lines and noted in italics. The catalytic dyad D25/D25’ is shown for structural context.

**Figure 6:**
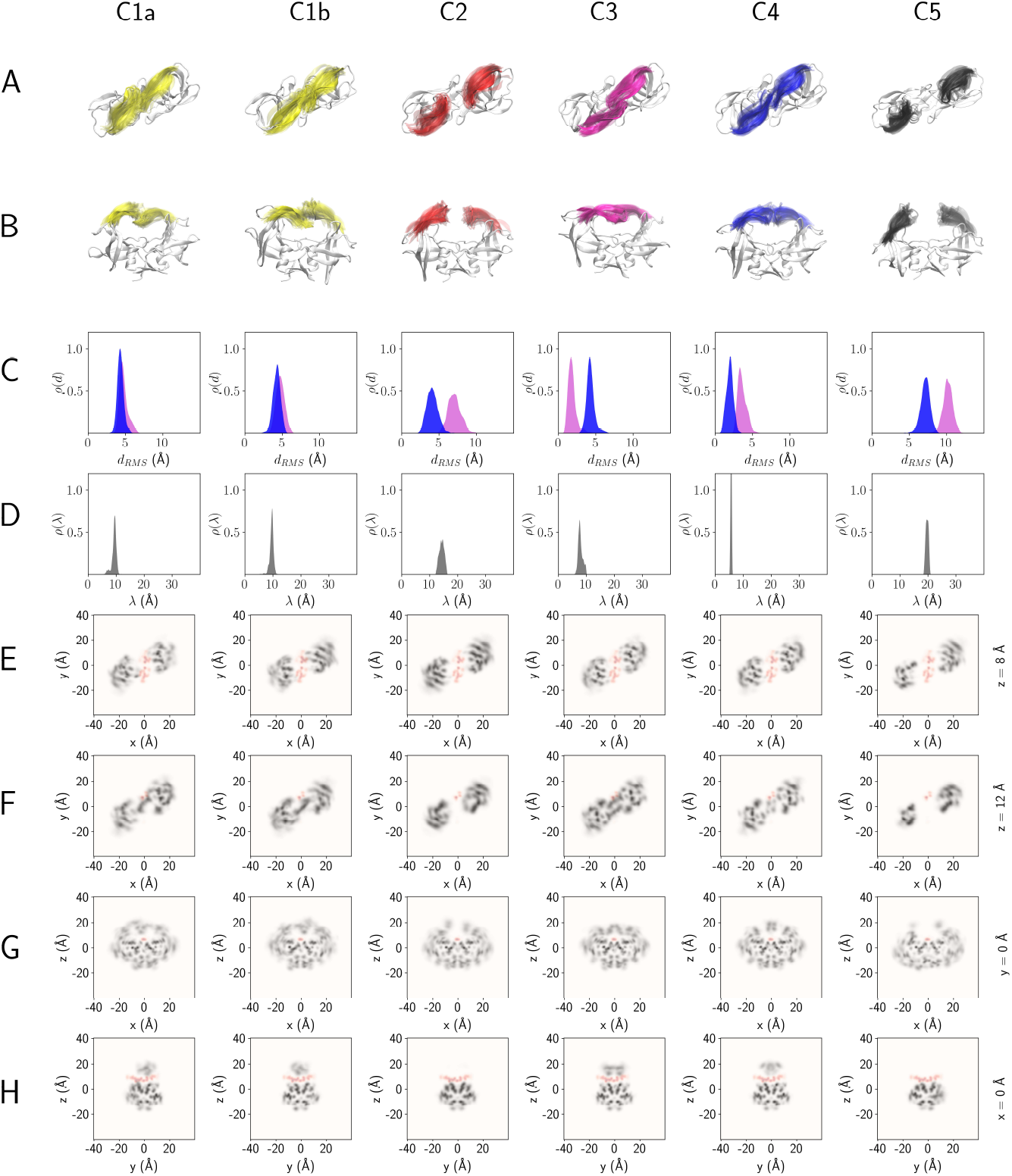
Structural characterization of representative conformers of each macrostate of HIV-1 apo-protease. (A) Top (x-y plane) and (B) side (x-z plane) views of ribbon representations of superimposed structures in each conformational ensemble, (C) Distribution (*ρ*(*d*)) of rootmean-square deviations (*d_rms_*) of C_*α*_ atoms of protease flaps (residues 43-58 in each monomer) with respect to crystallographic reference closed (pink) and semi-open (blue) structures. (D) Distribution of (*ρ*(λ)), the absolute flap tip separation distance (λ). (E-H) Cross-sections of superimposed volumetric density between the SP1-NC natural peptide in a closed-bound reference structure (red) and representative conformational ensembles of each macrostate (grey) in the x-y plane at z =8 Å (E) and z = 12 Å (F), the x-z plane at y = 0 Å (G) and the y-z plane at x = 0 Å (H).

Observations of C3 and C4 are further confirmed by unimodal distributions of flap-RMSD with respect to crystal-structures of closed and semi-open conformations for which C3 and C4 exhibit the smallest mean RMSDs, 1.8±0.5 Å and 2.0±0.4 Å, respectively (Figure 6(C)). All other macrostates exhibit significantly larger mean RMSDs with respect to both crystal structures. Analysis of the distribution of absolute flap tip separation distances (λ) shows that C3 and C4 all exhibit peak distances substantially lower than 10 Å, whilst C1a and C1b both peak at ~10 Å(Figure 6(D)). By contrast, C2 and C5 exhibit a relatively broad distribution of distances peaking at 15 Å and 20 Å respectively, implying that only these two states provide sufficient access to peptidomimetic ligands binding in the active site. The λ-distributions of all conformers for each of the top microstates of every macrostate exhibit very similar profiles respectively to the macrostate conformer sets selected here (Supporting Figure S2).

Cross-sections of averaged weighted atomic density over the conformers of each set provide further insight into the steric active-site accessibility of the corresponding macrostates (Figure 6(E)-(H)). The averaged density of a closed bound reference complex of HIV-1 protease with the SP1-NC cleavage peptide was calculated based on previous simulations.^117,118^ Superimposing densities calculated individually for the apo-protease (gray) and ligand (red) from this complex are consistent with both a snugness of fit and the inaccessibility of binding the enzymatically viable closed conformation (Supporting Figure S3). Furthermore, the comparable density profiles of the protein between the C3 conformation in the apo-protease simulations and the ligand-bound reference complex (Supporting Figure S3) further confirm that C3 corresponds to the closed conformation of the protease in the presence of bound ligand. The ligand exhibits significant density between −5 Å < *x* < 5Å, −15 Å < *y* < 15Å, 5 Å < *z* <10 Å. To provide corresponding steric context to the macrostates, the density of the peptide from this analysis was superimposed with those calculated from each of the representative conformer sets. Cross-sections in the x-y plane reveal that at z = 8 Å, states C1a, C1b and C3 exhibit partial occlusion at the edges of the active site, whilst states C2, C4 and C5 exhibit a zero-density lateral exit pathway in the y-direction. Moreover, states C2-C5 exhibit a zero-density groove corresponding to the active site that can accommodate the reference peptide density. However, by z = 12 Å, only macrostates C2 and C5 maintain this groove, whilst the flap extensions in the other conformations incur protein density that closes off peptide escape in the z-direction. Lateral cross-sections in the x-z and y-z planes confirm that only the C2 and C5 macrostates maintain a zero-density pathway out of the active site along the z-direction. Thus, combining exit pathways from the y- and z-directions, only macrostates C2, C4 and C5, corresponding to open, semi-open and wide-open conformations, respectively are selected as ligand accessible states for rigid body binding simulations.

### Gated ligand binding of peptidomimetic inhibitors

A set of peptidomimetic ligands (Figure 7), covering a range of experimentally determined association rate constants, *k_on_* ~10^4^ - 10^10^ M^−1^s^−1^, were chosen for Brownian dynamics simulations to each of the three chosen (C2, C4 and C5) accessible states of HIV-1 protease. The closest 100 conformers to the centroid of the most probable microstate were extracted for each of the accessible states. All three sets of conformers exhibit similar translational diffusion constants (< *D_T_*(*C*2) > = 9.90 ± 0.08 × 10^−11^ *m*^2^/s, < *D_T_*(*C*4) > = 1.00 ± 0.01 × 10^−10^ *m*^2^/s), < *D_T_*(*C*5) > = 9.80 ± 0.08 × 10^−11^ *m*^2^/*s*) and similar radii of gyration (*r_g_*(*C*2) = 1.99 ± 0.03 × 10^−9^ *m, r_a_*(*C*4) = 1.91 ± 0.02 × 10^−9^ *m, r_g_*(*C*5) = 2.04 ± 0.02 × 10^−9^ *m*). Translation diffusion constants and radii of gyration for the ligands range between 2.88 - 4.58 × 10^−10^ *m*^2^/s and 4.43 - 7.83 × 10^−10^ *m*, respectively, yielding a characteristic diffusional timescale, *τ_d_*, that ranges between ~ 12 - 16 ns depending on the ligand. By contrast, based on the outcome of the MSM and using Equation 17, the characteristic protein conformational transition timescale, *τ_c_*, is computed to be almost an order of magnitude larger (*τ_c_* = 85 ns), suggesting a ‘slow’ gating regime (*τ_c_* >> *τ_d_*). Thus, there is a kinetic gating factor to binding and via Equation 11, this is computed to be *γ* = 0.75.

**Figure 7:**
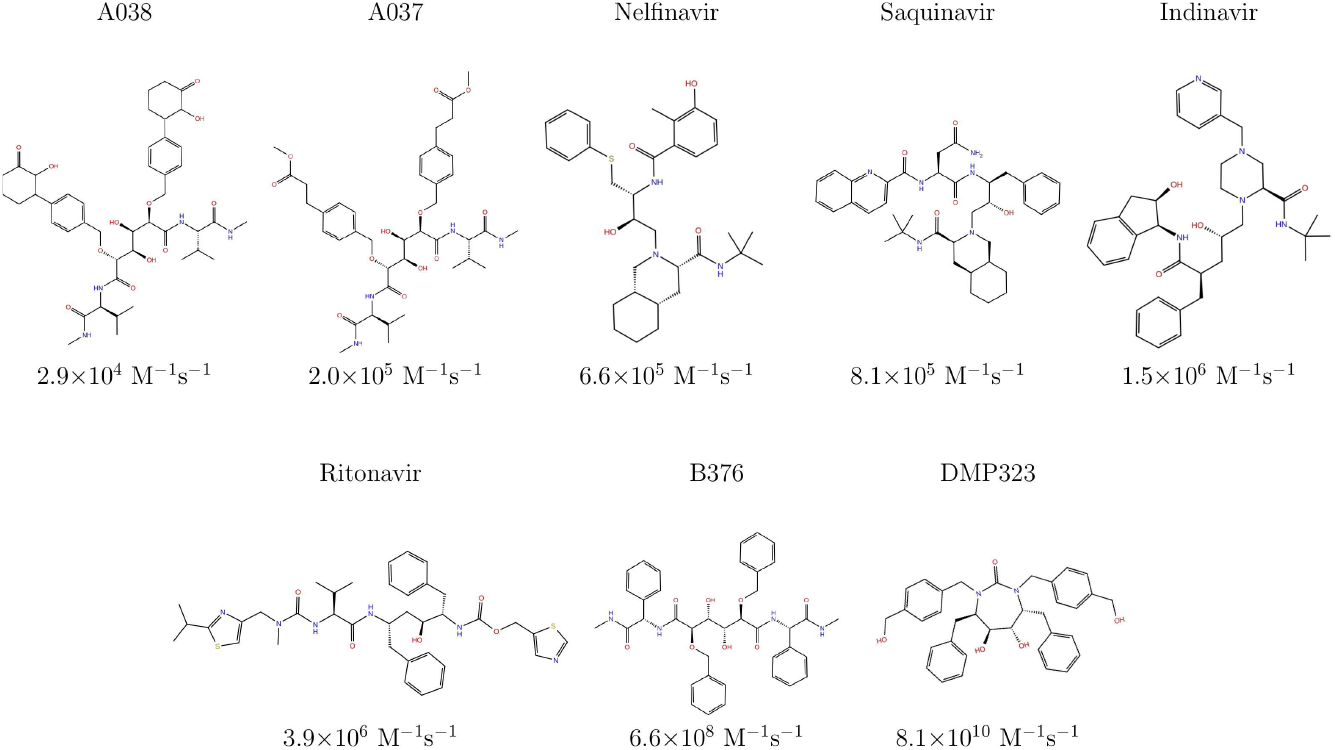
Chemical structures of the set of peptidomimetic ligands for which association rate constants were computed and which have a wide range of experimentally determined association rate constants, 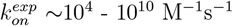.

Ungated rate constants 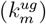 were then computed from BD simulations to each of the macrostates *m*=C2,C4 and C5, where the value for each state was computed using ensemble averaging over the set of 100 conformers that represented it. For each state, a number of complexes were formed at the reaction criteria corresponding to the bound state (see Methods). BD docking simulations were performed for each of the ligands binding to each of the accessible protein macrostates under these reaction criteria and the most popular clusters inspected. These confirmed that fulfillment of the reaction criteria corresponds to ligands binding within the active site of the protease (Supporting Figure S4). Similarly, BD simulations were carried out for ligands binding to the inaccessible states (C1 and C3). Lack of complexes formed within the active site of these two states thus further confirmed assignment of them as inaccessible (Supporting Figure S4).

Values for the ungated rate constants 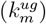 were combined to give the overall ungated rate constants for the set of accessible states (k^*ug*^) and finally multiplied by the gating factor *γ* to obtain the gated rate constants (k^*g*^). Our calculations show, as expected, that binding to the C4 (semi-open) state is slower than to the C2 (open) state and also slower than binding to the C5 (wide-open) state for all ligands except Nelfinavir and Saquinavir (Table 1). Furthermore, binding for most ligands is faster to C2 than C5, occasionally by severalfold, whilst for Ritonavir and B376 it is only marginally slower. The resulting combined ungated rate constants (k^*ug*^) are almost all within one order of magnitude of each other,ranging from *k^ug^*~60-750 *μ*M^−1^s^−1^ and, as the gating factor is 0.75, the gated rate constant range is only marginally reduced (*k^g^*~45-565 *μ*M^−1^s^−1^). This results in *k^g^* being up to several orders faster than the corresponding experimental association rate constants 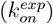 for most ligands, similar for B376, and several orders of magnitude slower for the fastest binder, DMP323 (Table 1).

**Table 1:** Comparison of experimental association rate constants 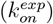 with computed ungated constants for individual 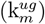 and multiple-combined (k^*ug*^) accessible states and corresponding gated rate constants (k^*g*^), computed with a gating factor of *γ* = 0.75. All rate constants are given in units of *μ*M^−1^ s^−1^.

| Ligand | $k_{C2}^{ug}$ | $k_{C4}^{ug}$ | $k_{C5}^{ug}$ | $k^{ug}$ | $k^g$ | $k_{on}^{exp}$ |
| --- | --- | --- | --- | --- | --- | --- |
| A038 | $417.00 \pm 5.24$ | $103.72 \pm 1.50$ | $330.67 \pm 5.70$ | $292.48 \pm 4.27$ | $219.36 \pm 3.20$ | $0.03 \pm 0.00$ |
| A037 | $349.53 \pm 2.81$ | $89.69 \pm 2.76$ | $262.07 \pm 5.66$ | $240.71 \pm 3.79$ | $180.53 \pm 2.84$ | $0.20 \pm 0.04$ |
| Nelfinavir | $156.96 \pm 3.70$ | $84.54 \pm 2.58$ | $61.77 \pm 0.80$ | $101.89 \pm 2.35$ | $76.42 \pm 1.76$ | $0.66 \pm 0.30$ |
| Saquinavir | $103.56 \pm 3.02$ | $48.59 \pm 3.21$ | $32.32 \pm 2.11$ | $62.11 \pm 2.76$ | $46.58 \pm 2.07$ | $0.82 \pm 0.16$ |
| Indinavir | $1052.30 \pm 8.50$ | $269.74 \pm 4.65$ | $866.02 \pm 10.31$ | $751.50 \pm 7.97$ | $563.63 \pm 5.98$ | $1.53 \pm 0.24$ |
| Ritonavir | $242.21 \pm 4.06$ | $39.90 \pm 1.28$ | $250.47 \pm 5.67$ | $184.16 \pm 3.79$ | $138.12 \pm 2.84$ | $3.92 \pm 0.11$ |
| B376 | $631.82 \pm 6.36$ | $112.97 \pm 2.71$ | $672.54 \pm 4.10$ | $489.77 \pm 4.47$ | $367.33 \pm 3.35$ | $205.00 \pm 187.00$ |
| DMP323 | $533.32 \pm 2.98$ | $157.72 \pm 2.26$ | $480.90 \pm 5.78$ | $401.87 \pm 3.74$ | $301.40 \pm 2.81$ | $25200.00 \pm 9990.00$ |

## Discussion and Conclusions

We have developed an approach to characterize the effects of gating by a multi-conformation protein consisting of subsets of accessible and inaccessible states on ligand association rate constants. Our approach first involves the construction of a kinetic network model of the apoprotein from atomistic MD simulations from which we identify meaningful conformations, compute their relative populations and interchange kinetics, structurally characterizing them in terms of ligand accessibility. We integrate the calculated first-order rate constants for conformational transitions into a multi-state gating theory from which we derive a characteristic timescale for conformational transitions (*τ_c_*) and a gating factor *γ*. This factor modulates the association rate constants of ungated ligand binding to the subset of accessible conformations (*k^ug^*) - which we compute here for a set of ligands from Brownian dynamics simulations - when *τ_c_* is significantly slower than the characteristic timescale for diffusional encounter (*τ_d_*) and enables us to quantify the degree of conformational gating by the protein.

Applied to HIV-1 protease, the kinetic network model of the apo-protein has macrostates corresponding to canonical crystallographic conformations - namely the well-characterized closed (C3) and semi-open (C4) states, as well as open (C2) and wide-open (C5) states previously reported from atomistic and coarse-grained simulations. It also reveals a previously unreported (C1) state (Figure 4). This latter conformation is an asymmetric doublet macrostate (C1a and C1b) in which the *β*-hairpin moiety, termed ‘flap’, of one monomer, reaches into and across the active site, making contacts with the opposite monomer and vice versa. These contacts are a mixture of those unique to the conformation (especially the flap tip I50 with the distal D30′), some only otherwise exhibited by the semi-open conformation, and others only otherwise by the closed conformation (Figure 5). Structural characterization reveals that only this state exhibits significant overlap in the averaged atomic density when superimposed with a putative natural substrate bound in the closed conformation. The other characterized states fit within an existing putative model of conformational change across the catalytic cycle^128^ - preferring a semi-open state in the apo protein, transitioning to an open/wide-open state during ligand encounter, then to the closed bound reactive state, and finally via an open state again for product release, back to the semi-open state. Our results suggest that C1 sterically partitions the active site, thus we term it ‘parted’ here. It may therefore assist in separating post-cleavage peptide fragments that are no longer covalently connected - and thus be favoured during product release, especially as it lies structurally on the pathway between closed and semi-open. CGMD simulations have suggested product release can occur even in the closed conformation.^129^ The parted conformation is kinetically disjoint from all other states except the semi-open in our apo-protein model - future atomistic simulations may reveal whether transition to the parted conformation from closed becomes kinetically favourable during product release.

Our kinetic model suggests that, individually, all of the macrostate conformations are almost thermodynamically indistinguishable from each other. The stationary distribution favours the subset of accessible (semi-open, open and wide-open) states, but even combined they are only ~1 kcal/mol more favourable than the sum of inaccessible states (parted and closed). Furthermore, there is comparatively rapid interchange between semi-open, open and wide-open states on the sub-*μ*s timescale whilst being an order of magnitude slower from these states to either parted or closed conformations. Interestingly, this results in a slow gating regime for ligand binding (*τ_c_* >> *τ_d_*), but because the calculated gating factor is not much smaller than unity (γ = 0.75), this can still be interpreted as the protease almost always being accessible or open to binding. This view broadly agrees with our previous study^95^ but here, we can quantify the estimated timescale of transition from semi-open to wide-open to be ~500 ns, in good agreement with a previous kinetic model.^96^ However, our model also suggests an equistable population for the wide-open conformation compared to the other conformations, rather than a rarely accessed state. The role of the wide-open state is still ambiguous as ligand binding does not seem to require it. Indeed, we find binding of almost all ligands studied here is faster to the open than to the wide-open state (Table 1). The relative stability of the wide-open state may instead be compatible with alternate roles. It has been observed that the protease makes direct non-specific interactions with the viral RNA - dramatically enhancing catalytic activity.^130^ Whilst the mechanism for this remains unclear, the wide-open state may further modulate RNA binding and thus indirectly regulate catalysis. The immature HIV-1 ribonucleoprotein forms a phase-separated condensate that allows influx of protease for its subsequent maturation.^131^ The conformational flexibility of the protease, including its ability to form the wide-open state may aid its mobility in such a crowded environment.

Possible slower conformational transitions that are not observed in our simulations may also play a role. Solution NMR experiments attributed a ~100*μ*s transition to a wide-open state.^89^ However, our model suggests that the transition to wide-opening occurs on the sub-*μ*s timescale. It is possible that the slower transition observed by NMR may be to yet another state. If such a state was inaccessible to ligand binding then it would significantly reduce the gating factor, and in turn reduce the gated association rate constant for all ligands.

The approach reported here enables us to dissect the potential contribution of conformational gating to a set of accessible states during the process of ligand binding and can be used to suggest possible binding mechanisms. For bi-conformational systems where the active and inactive conformations correspond to accessible (open) and inaccessible (closed) states, respectively, gating can be a modulating factor to the conformational selection (CS) mechanism. However, the complexity of HIV-1 protease draws out an important distinction between accessibility and active state. Even though a small ligand could bind directly to the active conformation (closed), this is largely inaccessible to peptidomimetic ligand binding, precluding a pure CS mechanism. Conformational gating therefore modulates accessible but intermediate binding states. For a pure induced fit (IF) mechanism, the ligand would need to initially bind a single conformation-gated accessible state, followed by a transition into the active (closed) conformation that may or may not be rate-limiting. However, several alternate complex binding mechanisms consistent with a dock and coalesce (DC) model may be possible, some of which are not addressed by our model.

In this study, we intrinsically decouple gating kinetics from ligand approach. However, proximity of a ligand to the protease may affect flap gating kinetics through transient intermolecular interactions. Therefore, even though the calculated gating factor suggests the protease active site is mostly accessible to binding, proximal ligand approach and/or loose binding away from the active site may alter this factor. Secondly, changes in protein and ligand conformations upon interaction are not accounted for in our rigid-body BD simulations. Transient interactions that involve flexible adaptation of a ligand to the protein along the binding pathway may result in either over- or underestimation of the ungated rate constant. Moreover, such interactions may allow a reactive flux through binding pathways different to first binding an accessible conformation. For example, a ligand binding loosely to the lateral exterior of the active site in a closed conformation, may kinetically trap the protease so that it cannot readily transition to an open state. However, this may be followed by partial opening or relaxation of the flaps to enable threading of the ligand into the active site to adopt a native bound state, with the rate constant depending on the context-specific flexibility of both the ligand and the protease and the ability to form transient interactions along the binding pathway. This deviation from pure CS, through a set of DC steps has been observed as a minor reactive flux in previous MD simulations of immature HIV-1 protease self-association.^53^ Conversely, initial encounters distal from the binding site may rapidly direct the ligand towards the active site, possibly by lower-dimensional diffusion across the protease surface. Similarly, another rationale for the existence of states such as the parted and wide-open may be to facilitate this rapid binding mechanism.

Here, we studied a set of inhibitors of HIV-1 protease whose association rate constants measured by surface plasmon resonance (SPR) vary across a wide range: 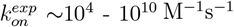. However, the calculated gated rate constants (*k^g^*) all lie within a narrow range from ~0.5-5.7 × 10^8^ M^−1^s^−1^. Only one inhibitor (B376), a small peptidomimetic ligand, exhibits a computed gated rate constant similar to the experimental association rate constant. The dominant binding mechanism for this inhibitor and those with a similar experimental rate constant is likely to one where binding is rate-limited by the initial conformation-gated diffusion into the active site and the subsequent transition to the native bound state is very fast.

Most inhibitors studied here are large peptidomimetic ligands (A038, A037, nelfinavir, saquinavir, indinavir and ritonavir) for which 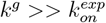 (Table 1) but for which 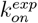 varies by only ~100-fold. These ligands are between ~100-10000 fold slower in binding than B376. Larger peptidomimetic chains have a greater degree of conformational freedom than smaller ones which is likely to be a dominant factor in this observed decrease in association rate constant. Their association may be limited by the gating of the conformational changes of the ligand or by subsequent protein-ligand changes to to the native bound configuration, both of which are not considered here. This mechanism is compatible with earlier findings from CGMD simulations on ligand binding. ^98^ However, it is possible that these ligands also exhibit the above-mentioned transient induced interactions not captured by our approach that may impede binding either by affecting the gating factor, ungated rate constant and/or by an alternate mechanism. Differential combinations of these contributions may explain the rate constant variation within this set of compounds.

Finally, the inhibitor DMP323, a smaller and more rigid cyclic urea compound, exhibits 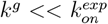 and is therefore not limited by gating of the diffusional association. Indeed, its rate constant exceeds the Smoluchowski diffusion limit. It is possible this fast rate constant could be due to an initial first encounter distal from the binding site followed by rapid surface diffusion to a final bound state. Previous MD simulations of a similar rapid cyclic urea binder with a 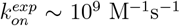 have suggested a binding mechanism where the ligand associates to the semi-open conformation and then induces an open conformation.^99,100^ However, it is also possible that the DMP323 measurement is an experimental artifact due to upper limits for reliable association rate constant measurements by SPR.^97^

Although, HIV-1 protease ligands likely exhibit several binding mechanisms, other proteins with gated binding sites may be largely modulated by the gating itself. In such cases, our approach may extend beyond suggested classification of different binding mechanisms and allow the computation of association rate constants that correlate with the overall experimental rate constants for different inhibitors. Finally, because our method only requires computing the apo-protein kinetics once from atomistic simulations and then using the extracted conformations for simulating the binding of each ligand using computationally inexpensive independent Brownian dynamics simulations that can be run in parallel, it could be effectively scaled for such protein systems for high-throughput ligand ranking.

## Acknowledgement

The authors thank Dr. Neil Bruce for assistance with SDA and discussions on the computation of rate constants, Dr. Gaurav Ganotra and Dr. Ariane Nunes-Alves for assistance in parameterizing ligand partial charges and diffusion coeffiecients, respectively, Dr. Ricard Argelaguet and Dr. Stefan Richter for technical assistance, Prof. Gianni De Fabritiis for sharing previous simulation trajectories from the GPUGRID project and Dr. Toni Giorgino, Prof. Gilles Mirambeau, Prof. Andreas Meyerhans and Prof. Ulrich Schwarz for discussions. This work was supported by the EU/EFPIA Innovative Medicines Initiative (IMI) Joint Undertaking, K4DD (grant no. 115366 to R.C.W.), the European Union Horizon 2020 Framework Programme (https://ec.europa.eu/programmes/horizon2020/en) for Research and Innovation under the Specific Grant Agreement no. 720270 (Human Brain Project SGA1, to R.C.W.), the Volkswagen Foundation “Experiment” Funding Initiative” (grant no. 93874 to S.K.S.), the amfAR Mathilde Krim Fellowship in Basic Biomedical Research (grant no. 108680 to S.K.S.) and the Klaus Tschira Foundation.

## Supporting Information Available

**Figure S1.** (A) Generalized Chapman-Kolmogorov test and (B) Distribution of microstate observation probabilities (χ) projected onto the λ-space.

**Table S1.** Top five most likely microstates that represent each macrostate.

**Figure S2.** Centroids of most probable microstate of each macrostate in the λ-space.

**Table S2.** Hydrogen bonds between flap and alternate monomers.

**Figure S3.** Cross sections of superimposed volumetric density of natural peptide and protein from a complex of HIV-1 protease.

**Table S3.** Sidechain-sidechain contacts between flap and alternate monomers.

**Figure S4.** Representative docked structures of peptidomimetic ligands to macrostates of HIV-1 protease.

**Data.** Simulation trajectories and output data pertaining to the results of the manuscript together with scripts for result generation and further analysis can be found on Zenodo at DOI: 10.5281/zenodo.5006612. Additional analysis scripts related to this data set can be found on Github at: https://github.com/kashifsadiq/hiv1pr-msm/.

## Notes

### Competing Interest Statement

The authors have declared no competing interest.

### Summary of Updates

Introduction section text revised Discussion and Conclusions section text revised Figure 2 revised

