## Supporting Information for "A multiscale approach for computing gated ligand binding from molecular dynamics and Brownian dynamics simulations"

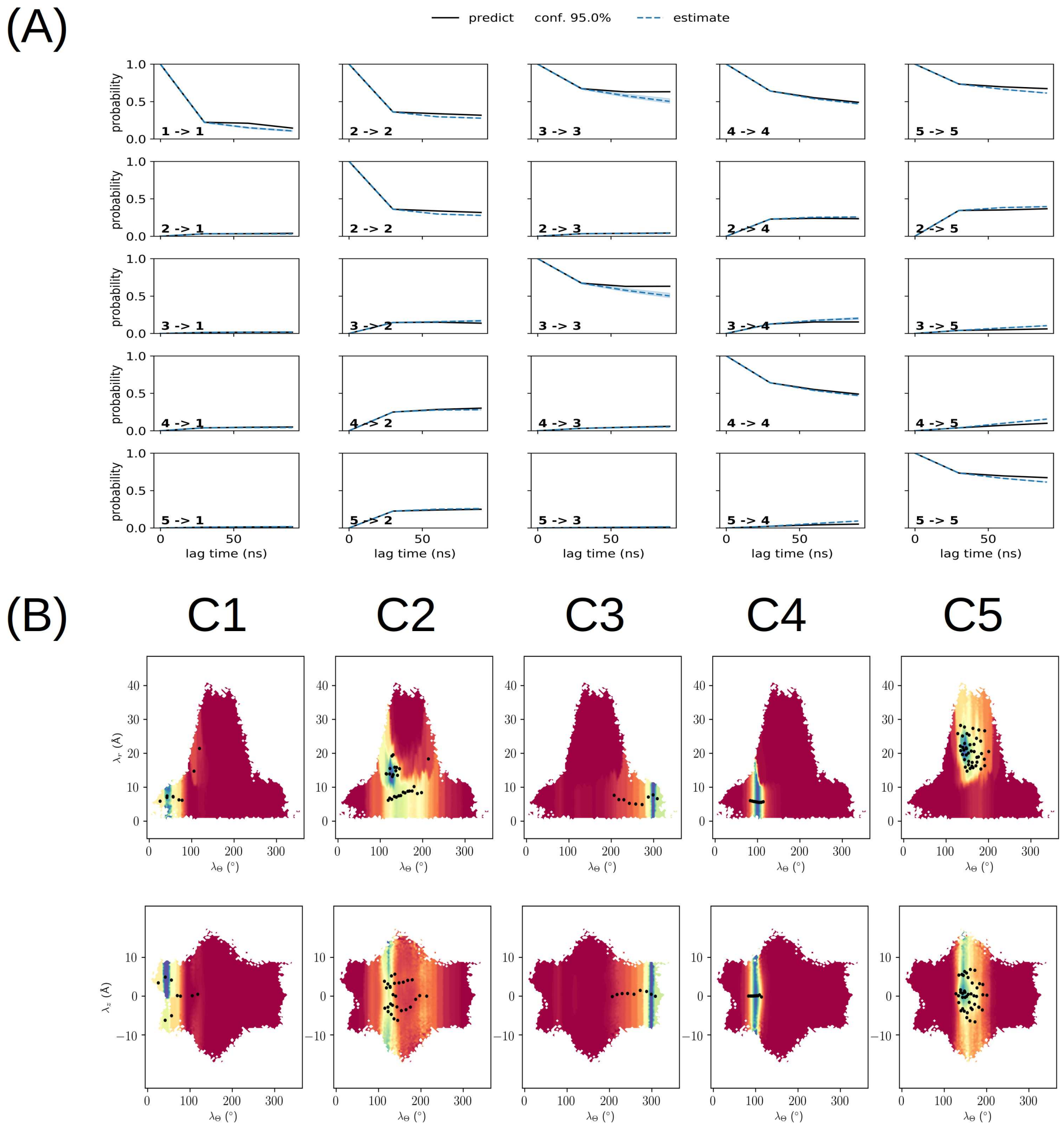


Figure 1: (A) Generalized Chapman-Kolmogorov test of macrostate probability decay for an MSM constructed at $\tau$=30 ns with five macrostates. Good agreement is observed between the predicted probability decay from the model ($p^k_{IJ}(\tau)$) and the expected decay from MSMs constructed at higher lag times ($p_{IJ}(k\tau)$). (B) Distribution of probabilities ($\chi$) of five kinetically distinct coarse-grained macrostates across all microstates of the MSM, projected onto the $\lambda$-space. Microstate distributions closely match the regions of the $\lambda$-space characteristic of conformations C1-C5 and their corresponding microstate centroids (black dots). Probabilities are color-coded from lower (red) to higher (blue).

Table 1: Top five most likely microstates that represent each macrostate. The top microstate for each corresponding macrostate is highlighted in bold.

| Macrostate (I) | Microstate ID (j) | Observation probability ($\chi_{I,j}$) |
|---|---|---|
| C1a | **79** | 0.229 |
| C1b | **12** | 0.133 |
| C1 | 56 | 0.12 |
| C1 | 53 | 0.104 |
| C1 | 94 | 0.093 |
| C2 | **98** | 0.045 |
| C2 | 77 | 0.042 |
| C2 | 26 | 0.035 |
| C2 | 80 | 0.031 |
| C2 | 7 | 0.031 |
| C3 | **2** | 0.278 |
| C3 | 40 | 0.177 |
| C3 | 32 | 0.146 |
| C3 | 19 | 0.08 |
| C3 | 67 | 0.062 |
| C4 | **1** | 0.135 |
| C4 | 78 | 0.133 |
| C4 | 76 | 0.122 |
| C4 | 39 | 0.118 |
| C4 | 14 | 0.096 |
| C5 | **6** | 0.039 |
| C5 | 48 | 0.037 |
| C5 | 47 | 0.035 |
| C5 | 55 | 0.035 |
| C5 | 87 | 0.034 |

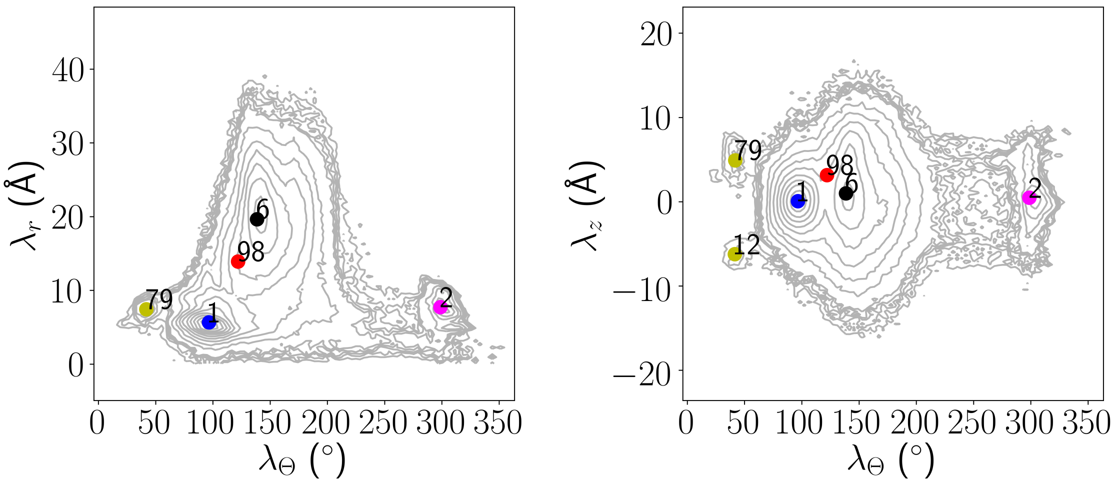


Figure 2: Centroid of the most probable microstate for each macrostate (C1: yellow, C2: red, C3: magenta, C4: blue, C5 black) superimposed on 2D projections of the 3D-$\lambda$ space. For C1, the top two most probable microstates are shown corresponding to sub-states C1a (79) and C1b (12), respectively.

Table 2: Hydrogen bonds between flap and opposite monomers.

| Macrostate | Hydrogen bond | $f_{occ}$ | $d_{ha}$ (Å) | $\theta_{dha}$ (°) |
|---|---|---|---|---|
| C1a | G48′:N-H–O:I50 | 0.67 | 2.94 ± 0.15 | 163.39 ± 7.19 |
| C1b | G48:N-H–O:I50′ | 0.64 | 2.93 ± 0.15 | 162.93 ± 7.25 |
| C2 | - | - | - | - |
| C3 | - | - | - | - |
| C4 | G51:N-H–O:G49′ | 0.68 | 2.98 ± 0.16 | 163.21 ± 6.95 |
| | G51′:N-H–O:G49 | 0.69 | 2.97 ± 0.16 | 162.83 ± 6.99 |
| C5 | - | - | - | - |

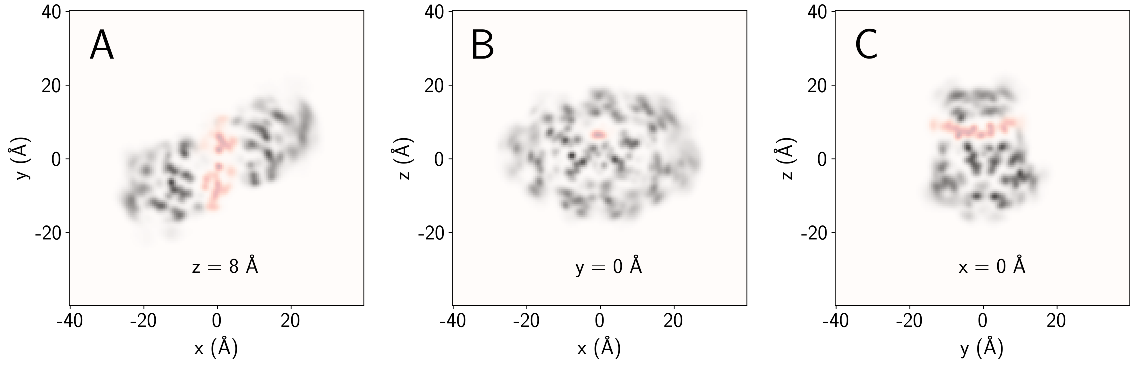


Figure 3: Cross sections of superimposed volumetric density in the (A) x-y, (B) x-z and (C) y-z planes, between the SP1-NC natural peptide and closed bound conformation of HIV-1 protease, both extracted from the same crystal bound reference complex (PDB: 1KJ7).

Table 3: Sidechain-sidechain contacts between flap and alternate monomers.

| Residue Pairs | | $d_{ss}$ (Å) | | | | | |
|---|---|---|---|---|---|---|---|
| Mon A | Mon B | C1a | C1b | C2 | C3 | C4 | C5 |
| D30 | I50′ | - | 4.7 ± 1.4 | - | - | - | - |
| I50 | D30′ | 4.7 ± 1.4 | - | - | - | - | - |
| V32 | I50′ | - | 4.6 ± 1.4 | - | 4.9 ± 1.1 | - | - |
| I50 | V32′ | 4.7 ± 1.5 | - | - | 4.7 ± 0.9 | - | - |
| M46 | I50′ | 4.3 ± 0.9 | - | - | - | - | - |
| I50 | M46′ | - | 4.6 ± 1.2 | - | - | - | - |
| I47 | I50′ | - | 4.1 ± 0.4 | - | 4.3 ± 0.6 | - | - |
| I50 | I47′ | 4.2 ± 0.4 | - | - | 4.2 ± 0.5 | - | - |
| G48 | G49′ | - | 4.6 ± 0.6 | - | - | - | - |
| G49 | G48′ | 4.6 ± 0.5 | - | - | - | - | - |
| G48 | I50′ | - | 4.6 ± 0.4 | - | - | 4.2 ± 0.4 | - |
| I50 | G48′ | 4.4 ± 0.4 | - | - | - | 4.2 ± 0.5 | - |
| G49 | I50′ | - | - | - | - | 4.5 ± 0.4 | - |
| I50 | G49′ | - | - | - | - | 4.5 ± 0.4 | - |
| G49 | F53′ | - | 3.9 ± 0.8 | - | - | - | - |
| F53 | G49′ | 4.1 ± 0.7 | - | - | - | - | - |
| I50 | I50′ | - | - | - | - | 4.8 ± 0.7 | - |
| I50 | F53′ | - | 3.9 ± 0.4 | - | - | 5.0 ± 0.9 | - |
| F53 | I50′ | 4.0 ± 0.5 | - | - | - | 4.9 ± 0.9 | - |
| I50 | I54′ | - | - | - | 4.3 ± 0.6 | - | - |
| I54 | I50′ | - | - | - | 4.2 ± 0.5 | - | - |
| I50 | T80′ | - | - | - | 4.2 ± 0.9 | - | - |
| T80 | I50′ | - | - | - | 4.2 ± 0.9 | - | - |
| I50 | I84′ | - | - | - | 5.0 ± 1.3 | - | - |
| I84 | I50′ | - | - | - | 4.9 ± 1.3 | - | - |
| G51 | G51′ | - | - | - | 4.7 ± 0.7 | 4.4 ± 0.9 | - |
| G51 | G52′ | - | - | - | 4.3 ± 0.9 | - | - |
| G52 | G51′ | - | - | - | 4.3 ± 0.9 | - | - |
| G51 | I54′ | - | - | - | 5.0 ± 0.6 | - | - |
| I54 | G51′ | - | - | - | 5.1 ± 0.6 | - | - |

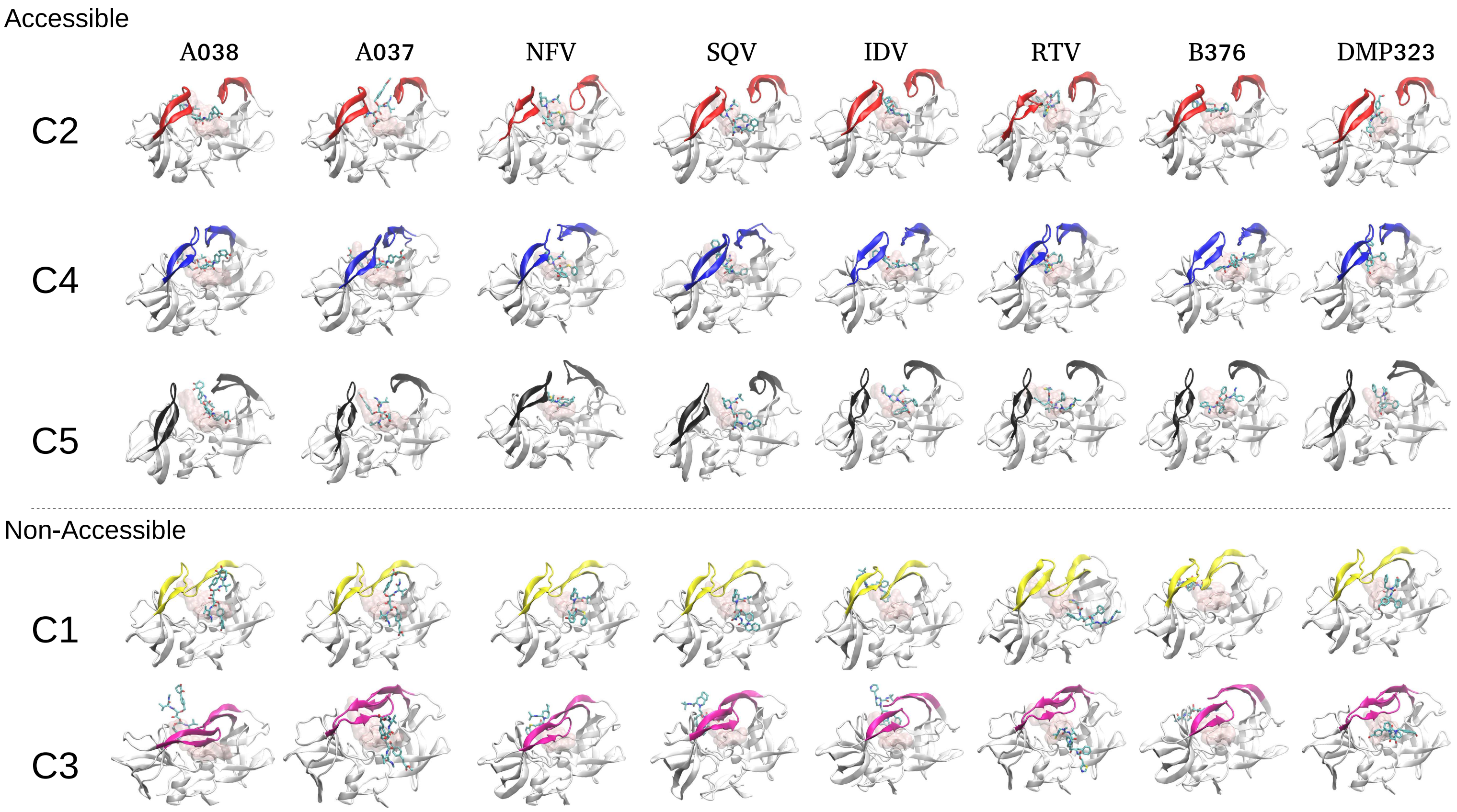


Figure 4: Representative docked structures of peptidomimetic ligands to macrostates of HIV-1 protease. Pink regions correspond to surface representations of the given ligand in the crystal structure of the complex, after alignment of residues 1-42 and 59-99 of each monomer of the protease.
